# Using insects for sustainable waste management of superabundant animals

**DOI:** 10.1101/2025.11.03.686388

**Authors:** Carlos López-Manzano, Hayat Mahdjoub, Luis Rodrigo Arce-Valdés, Rassim Khelifa

## Abstract

Management of fecal waste from superabundant wildlife in urban areas is a key environmental and public health concern, yet even in developed countries, sustainable solutions that implement circular economy principles are still lacking. We tested the potential of black soldier fly (BSF) larvae for valorizing organic waste of the widespread Canada goose: converting droppings into larval protein while yielding frass fertilizer. Larvae were able to survive, develop, and degrade the goose feces. For instance, larvae degraded 56% of goose feces compared to 63% of a control diet. Sterilization of feces reduced the performance of larvae. We tested the fertilization properties of the insect frass on an aquatic plant (duckweed) and we found growth enhancement of 32% at 10 g·L^-1^ compared to a standard Hoagland’s media. Our results provide insights into how to sustainably manage urban fecal waste from overabundant species while producing protein and a nutrient-rich fertilizer for plants.

## Introduction

Organic waste management, as a cornerstone of the global circular economy, plays a pivotal role in the transition toward sustainable and resilient urban systems ^1^. In 2014, the total mass of animal feces (primarily from cattle, chickens, and sheep) was estimated at 3.9□×□10^12^□kg per year—about four times that produced by humans—and is projected to reach at least 4.6□×□10^12^□kg by 2030 ^2^. Beyond these managed fecal streams, superabundant wild species such as birds and bats generate substantial quantities of fecal biomass that remain untreated, including in and around urban areas ^3,4^. This neglected input accumulates in domestic environments, where it poses significant risks for pollution, disease transmission, and economic costs of sanitation ^5,6^. Meeting the anticipated growth of urban populations—projected to reach 6.6 billion people by 2050 ^7^—will likely exacerbate these pressures, intensifying the need for innovative waste management strategies. Harnessing fecal biomass from superabundant species through circular solutions has the potential to sustainably transform waste into valuable resources^8^, thereby contributing to the broader global green transition ^9,10^.

The Canada goose (*Branta canadensis*), a common and iconic bird in North America, thrives in urban, suburban, and rural areas ^11^. Once depleted by overhunting and habitat loss, populations rebounded following conservation measures and landscape modifications ^12^. Their population grew by over 7% annually from 1966 to 2019 according to the North American Breeding Bird Survey ^13^, and burgeoning urban populations have created ecological, social, and health challenges ^14–18^. Large flocks of Canada geese can damage vegetation, disrupt agriculture, and outcompete other wetlands birds ^19,20^. Pathogens in their feces make parks, sports fields, and shorelines unsanitary ^21^. Each goose produces 1.5 pounds (0.68 kg) of feces daily, defecating every 20 min ^22^. Despite the scale of the issue, feces management remains poorly regulated in North America. Specialized companies collect goose feces, but disposal is often inadequate (e.g., disposing of it in landfills alongside regular waste) and does not involve recycling or composting. Similarly, feces-collection machines are commercially available (e.g., Tow and Collect, USA), yet no dedicated treatment or recycling strategies have been implemented. These limitations highlight the need for improved management solutions to mitigate human–wildlife conflicts associated with Canada geese.

Recently, the black soldier fly (*Hermetia illucens*; BSF) has gained increasing attention as a sustainable tool for biowaste management ^23,24^. Due to its wide trophic range, this insect can digest food scraps, agricultural byproducts, and livestock manure ^25^. BSF larvae efficiently convert waste into high-protein livestock feed ^26–28^. AAFCO allows dried whole BSF and protein meal for salmon, trout, and char aquaculture ^29^. Some US states have approved insect-based pet foods, while others await AAFCO and FDA guidance ^29^. NACIA advocates for North American insect industry. Despite its widespread use to degrade waste, the BSF has not been tested for managing superabundant wildlife like the Canada goose. This novel research avenue may offer a sustainable solution for Canada goose waste management.

An organic fertilizer made from BSF frass, a nutrient-rich byproduct of larval growth, is being studied as a sustainable alternative to synthetic fertilizers ^30^. BSF frass (excrement) is a promising soil amendment that boosts plant growth, crop yields, soil microbial activity, and nutrient availability due to its nitrogen, phosphorus, and potassium content ^31–33^. The larval feedstock determines efficacy ^32^. Frass produces biomass and uses nitrogen better than commercial organic fertilizers ^34^. Frass application research has mostly focused on terrestrial crops, leaving aquatic plant growth potential unexplored ^35^. It is unclear how nitrogen-rich aquatic bird frass ^36^ supports the cultivation of high-feed and food potential aquatic plants.

Lentic freshwater systems have duckweeds (Araceae: Lemnoideae) ^37^. They are the smallest, fastest-growing plants that can reproduce vegetatively in ideal light and temperature ^38^. Their 1.34–4.54-day doubling times demonstrate their impressive growth potential ^38^. Natural duckweeds feed herbivores, shelter macroinvertebrates, and reduce nutrient loads ^39^. They are ideal ecotoxicology, ecology, and evolution model organisms due to their biology. Duckweeds are increasingly important for sustainable industries beyond their ecological value. Wastewater treatment, livestock feed, human food, and biofuel production have been studied ^40^. Rapid duckweed growth, high protein content, and ability to thrive in nutrient-rich environments make them promising for environmental management and commercial applications ^41^. BSF frass as fertilizes supports *Lemna minor* growth ^35^, making it suitable for sustainable waste management. Little is known about its fertilizer value for duckweed.

In this study, we experimentally tested the effectiveness of black soldier fly larvae in degrading Canada goose feces and assessed the potential of the resulting frass for promoting the growth of an economically important aquatic plant (duckweed) (Fig. 1). Specifically, we first conducted a field survey in urban parks of Southern Quebec and Ontario (Canada) to assess whether larger flocks of the Canada goose are correlated with larger numbers of feces. We then tested whether the Canada goose feces are consumed and degraded by BSF larvae using a laboratory experiment that compares the performance of the insect in a control diet (Gainesville) with different treatments with increasing proportions of feces. Furthermore, to assess the potential of BSF frass as a fertilizer for cultivating duckweed, we evaluated the growth response of duckweed (*Lemna minor*) to different concentrations of fresh feces of the Canada goose and BSF frass.

**Figure 1.**
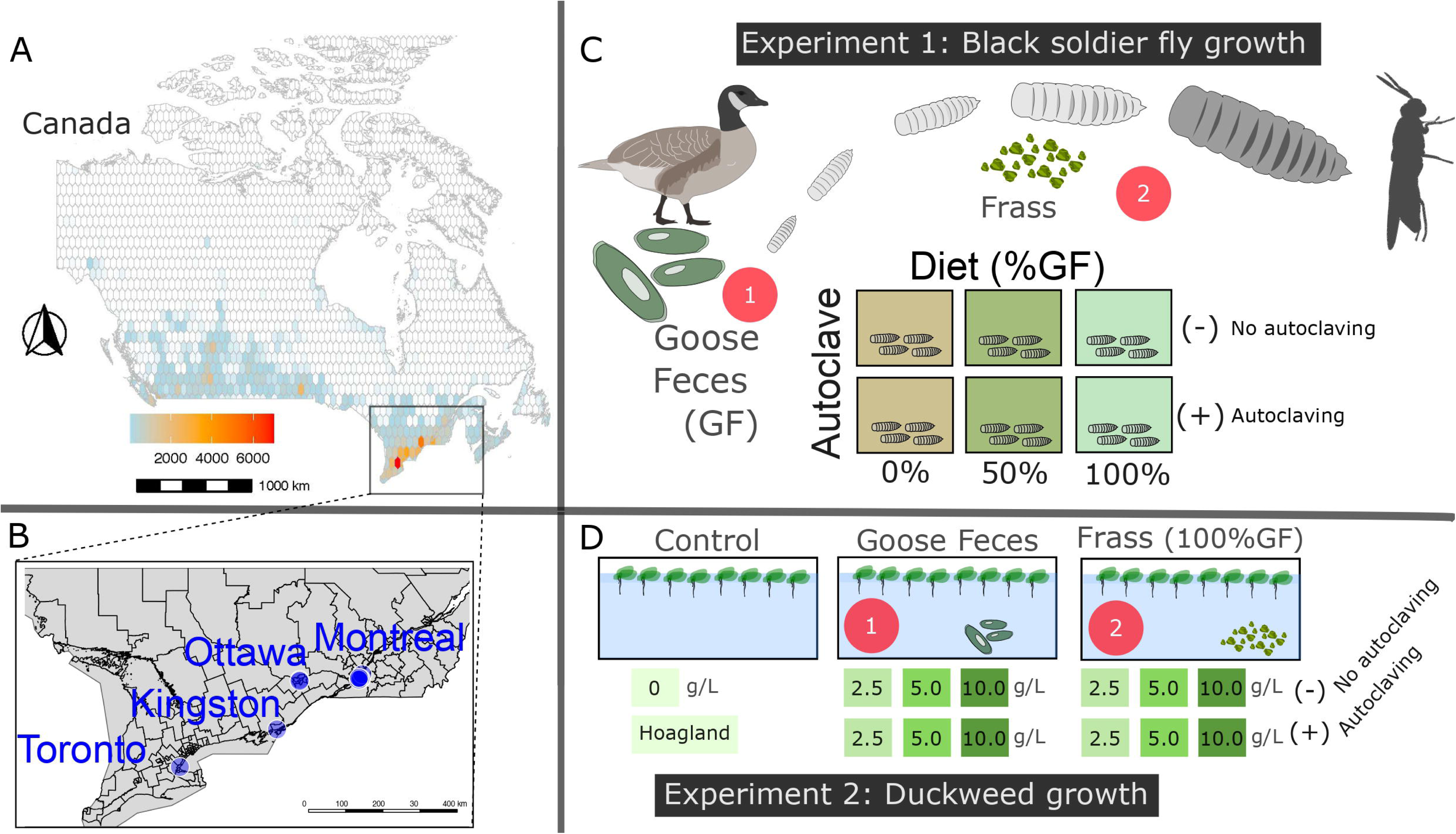
Field survey and experimental design. Geographic distribution (a) of the Canada goose in Canada. The map was based on occurrences reported on GBIF database (duplicate locations were removed). The gradient shows the number of reported sites where the Canada goose was observed. (b) Study sites for field survey. (c) Experimental framework (Experiment 1) to assess the potential of black soldier fly larvae in degrading the Canada goose droppings. (d) Experimental framework to assess the potential of fresh and processed (BSF frass) dropping of the Canada goose (Experiment 2).

## Results

### Canada goose field survey

The abundance of the Canada goose in the studied urban green spaces averaged 73.0±69.5 individuals (Table S1). The number of droppings recorded across our 100 m transect had an average of 54.9±34.7. We found a significant positive correlation between the number of Canada geese and the number of droppings (estimate = 0.46 ± 0.06 (SE), t=7.54, R^2^ = 0.85, P < 0.0001; Fig. 2), indicating that the amount of fecal material produced by the bird increased with the size of the flock. The fresh weight of the Canada goose single droppings averaged 13.51±3.30 g (N=12), and the dry weight represented 15% of the fresh weight.

**Figure 2.**
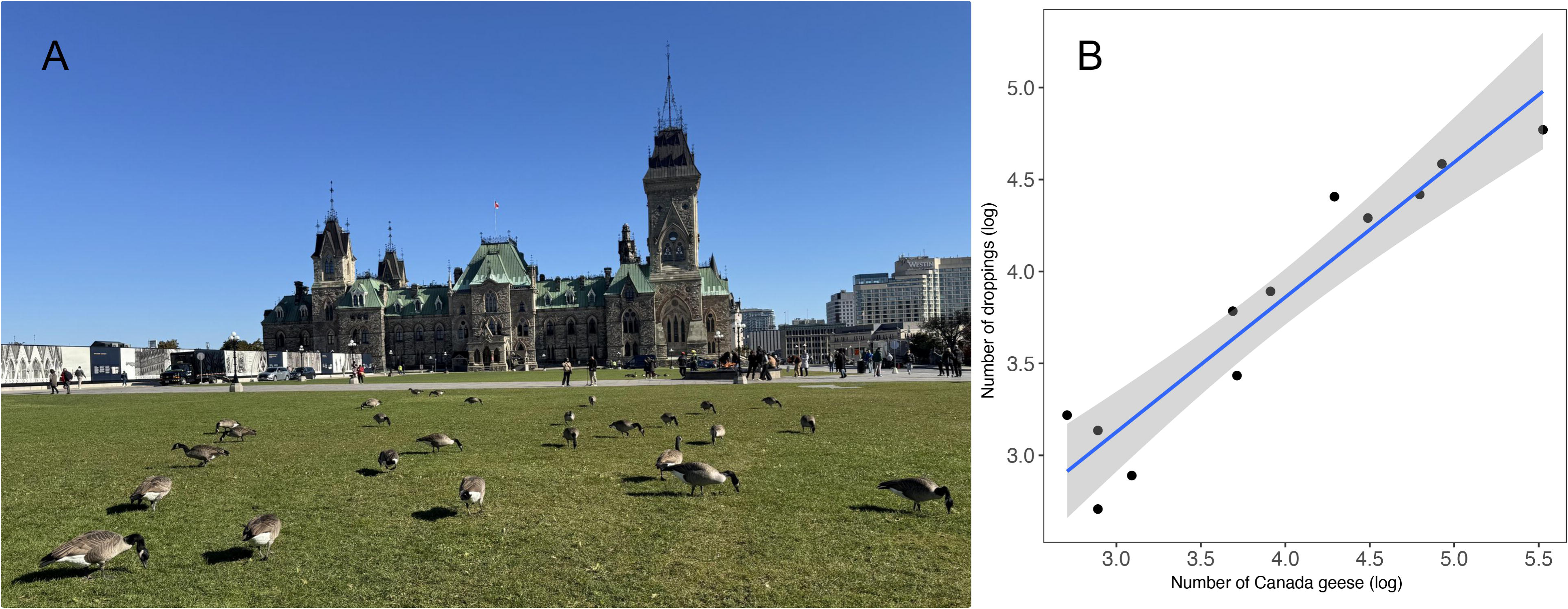
Canada geese (*Branta canadensis*) in urban ecosystems. **a,** Canada geese in an urban green space at Parliament Hill in Ottawa (ON, Canada). **b,** Correlation between the number of droppings and the number of Canada geese. Data collected in urban green spaces in Southeast Canada.

### BSF larvae performance on goose dropping

#### Larval growth rate

Larvae showed different larval growth across the different diets (Fig. 3a and Table S2). Larvae reared with the control diet (F0) and F50 showed faster growth rates than F100. During the first week of larval growth, the growth rate was faster in F50 than in F0 and F100, as revealed by the interaction between time and diet (F_2,528_=139.0, P<0.0001). The significant interaction between diet and sterilization (F_2,528_=12.6, P<0.0001) indicates that autoclaving the media had no effect on the control diet.

**Figure 3.**
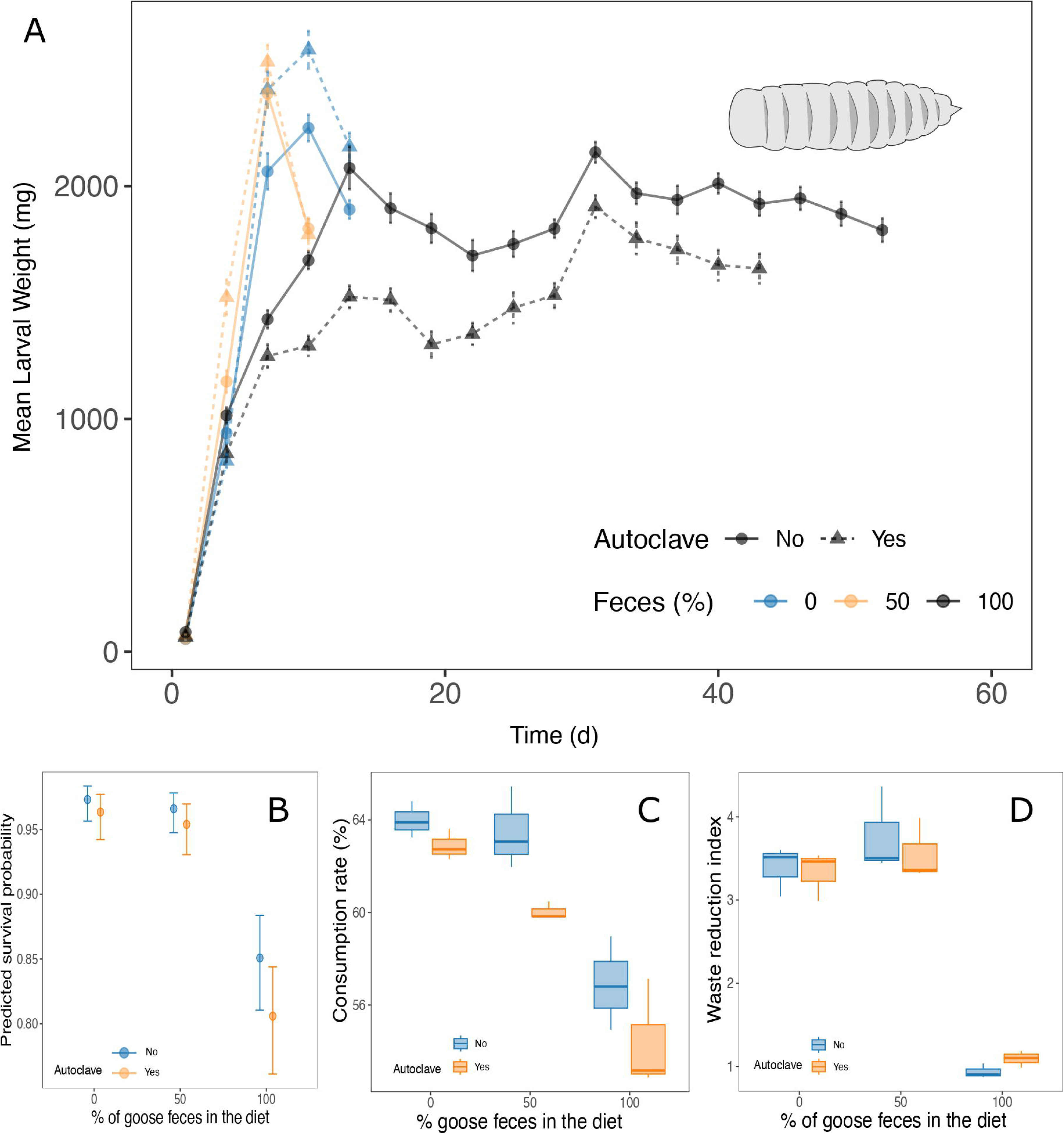
Larval performance in different diet treatments. **a,** Growth rate of black soldier fly (*Hermetia illucens*) larvae. BSF was reared in different diets with increasing proportions of Canada goose feces under two sterilization treatments. **b,** Survival probability of larvae in difference diets. **c,** Consumption rate of the media by BSF larvae. **d,** Waste reduction index of BSF reared in different diet treatments.

#### Larval survival

Larval survival was the highest in the control diet (F0; 96.4% [95% CI: 94.2– 97.7%]) and in the mixture diet (F50: 50% control with 50% feces; 95.4%, 95% CI: 93.1–97.0%). There was no significant difference between F0 and F50 (GLM: *z* = –0.78, *P* = 0.438). Larvae strictly growing on the feces-only diet (F100) had significantly lower survival (80.6%, 95% CI: 76.1–84.4%; *z* = –7.18, *P* < 0.001) compared with both F0 and F50. Sterilization (autoclaving) did not significantly affect survival (GLM: *z* = 1.77, *P* = 0.077) (Fig. 3b).

#### Consumption rate

The highest consumption rate (% dry-matter reduction) was observed in control treatments (64.0 ± 0.8% in raw media and 62.9 ± 0.7% in sterile media) (Fig. 3c). F100 treatments had the lowest consumption rates (56.9 ± 2.0% and 54.4 ± 2.4%), while F50 treatments had slightly lower values (63.5 ± 1.8 % and 60.0 ± 0.4 %). Fecal content and autoclaving had significant effects, but their interaction was not significant (P = 0.435; Table S3). Across all fecal levels, consumption in sterile media was significantly lower than raw media (P = 0.007, Table S4).

#### Waste Reduction Index (WRI)

Waste reduction was highest in F50 treatments (3.77 ± 0.52 and 3.56 ± 0.37 g DM d□^1^ in raw and sterile media, respectively), followed by controls (3.39 ± 0.30 and 3.33 ± 0.30 g DM d□^1^) (Fig. 3d). The F100 treatments had the lowest waste reduction rates (0.94 ± 0.09 g DM d□^1^ and 1.09 ± 0.10 g DM d□^1^). Fecal content had a significant effect (Table S10), while autoclaving does not and the interaction is not significant (P = 0.611; Table S5). At any fecal level, sterilization did not reduce WRI (P = 0.802: Table S6).

#### Development time and adult traits

Development time differed significantly across diet treatments (t=7.01, P<0.0001) (Table S7, Fig. 4a). Males had longer development times than females (t = 6.49, P < 0.0001). Adult dry mass declined sharply with increasing goose feces content and differed between all diet levels (F_2,103_ = 274.5, P < 0.0001) and was 23 % higher in females than in males (F_□,□□□_= 28.0, P < 0.0001) (Table S8, Fig. 4b). Adult lifespan was significantly different across diets (P = 0.0134, supplementary methods). Lifespan was significantly shorter in F100 compared to F0 (z=5.61, P <0.0001) and F50 (z=3.76, P =0.0005) (Fig. 4c).

**Figure 4.**
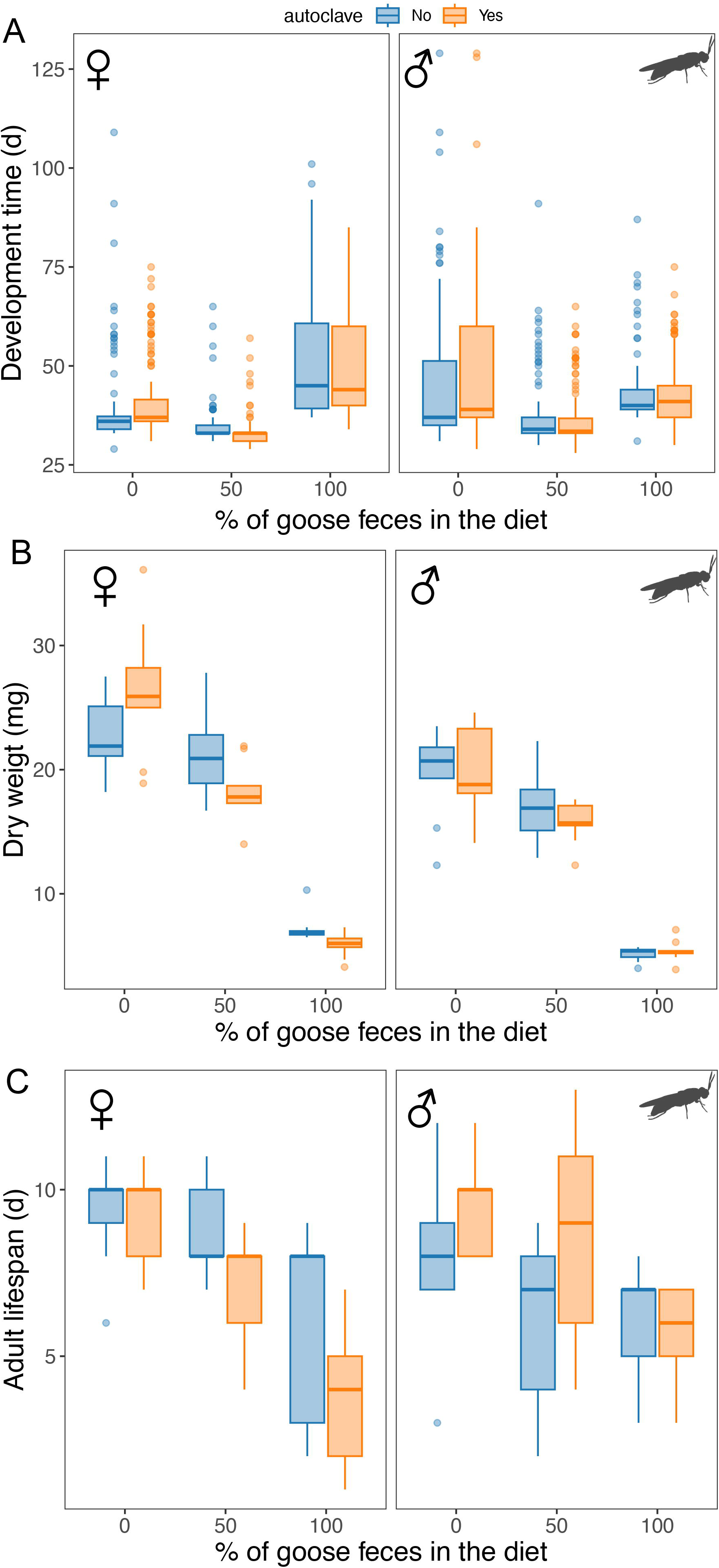
Larval development and adult traits of the black soldier flies (*Hermetia illucens*) reared in different diets with increasing proportions of Canada goose feces. a, Development time (egg hatching to adult). **b,** Adult dry body mass. **c,** Adult lifespan.

### Goose feces for duckweed growth

#### Duckweed growth rate

Frass treatments enabled greater and more sustained duckweed growth than feces or controls (Fig. 5a-c, Fig. S1, Table S9-11). Peak yields after 28 d increased with frass concentration: 5.32, 5.40 and 6.02 fronds at 2.5, 5 and 10 g·L□^1^, respectively (95 % CI ranges 4.99–6.37). Feces treatments plateaued lower (4.65–4.95 fronds). The 10 g·L□^1^ frass curve had a significantly later inflection point (4.83 days [3.41–6.25]), indicating prolonged, productive growth. Hoagland solution reached 4.57 fronds [4.38–4.77], still below high-frass. Autoclaving did not change the asymptote overall (4.72 vs 4.65; p = 0.19) but significantly delayed the inflection point by 2.02 days [1.23–2.81], slowing early growth. A significant three-way interaction (medium × concentration × autoclaving) on the asymptote revealed that sterilization raised frass yield at 5 and 10 g·L□^1^ yet lowered it at 2.5 g·L□^1^, whereas in feces it reduced yield at 2.5 and 10 g·L□^1^ and had no effect at 5 g·L□^1^ (Supplementary results).

**Figure 5.**
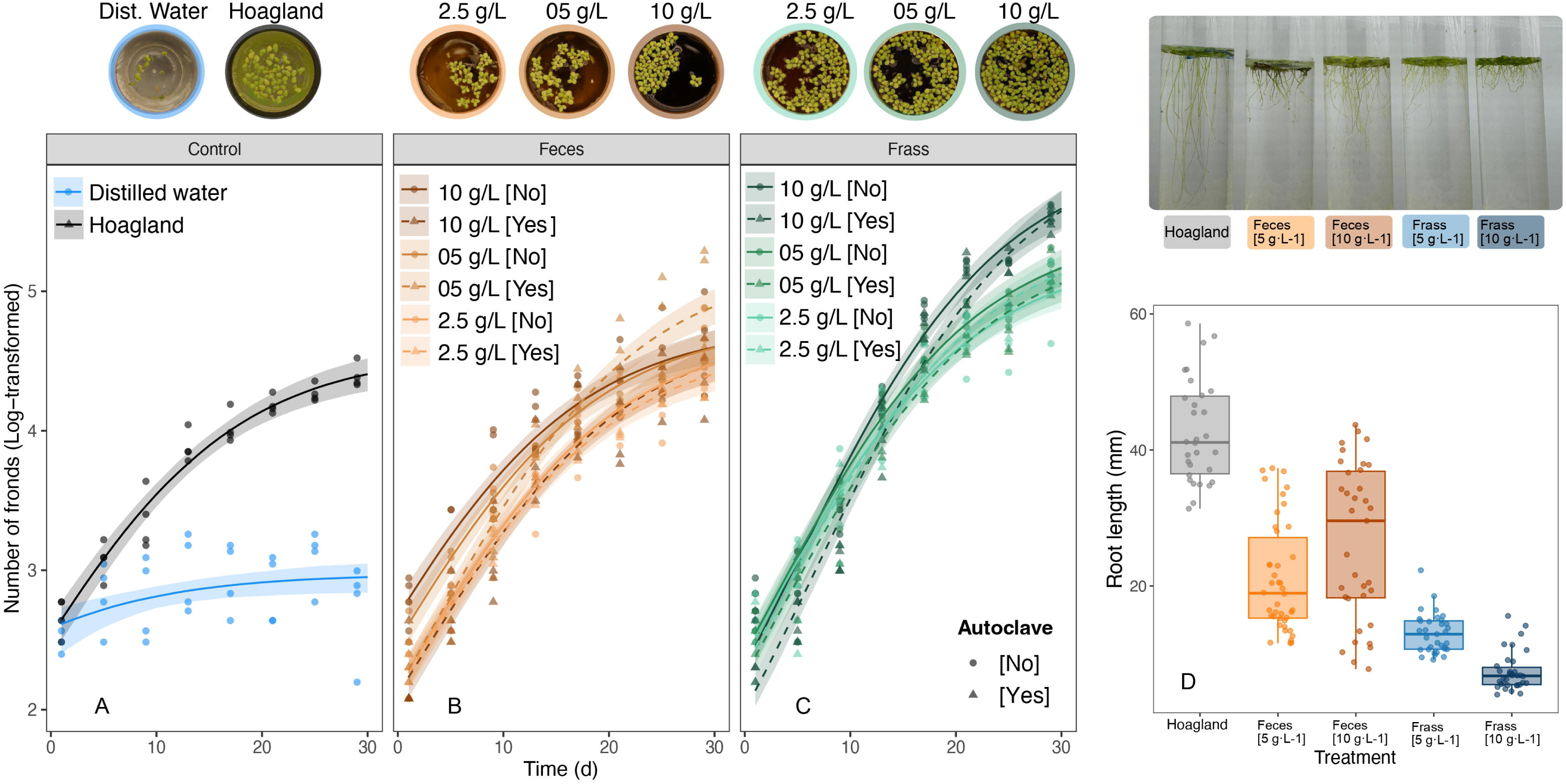
Growth curves of the duckweed (*Lemna minor*) in different media. **a,** control (distilled water and Hoagland’s media). Three concentrations (2.5, 5, 10 g·L^-1^) of the **b,** fresh goose feces and **c,** insect-based frass based on the processing of the Canada goose feces by black soldier fly larvae (*Hermetia illucens*). The two treatments (feces and frass) were prepared in the presence and absence of autoclaving. The images above the figure represent the state of duckweed growth after four weeks of growth. Note the difference in abundance of duckweed fronds. **d,** Root length after two weeks of growth in different frass-based media. The image above shows a sample of root length after four weeks of growth.

#### Root length

Root length was significantly shorter in the frass treatment than the feces and Hoagland’s media (F_4,169_= 106.4, P<0.0001; Tukey test: P<0.01) (Fig. 5D). Roots were on average 76% and 57% shorter in the frass and feces treatment than Hoagland’s, respectively. The largest reduction in root length was observed in the Frass 10 g·L^-1^ treatment, where roots were 83% shorter compared to those grown in Hoagland’s medium (Fig. 5d).

## Discussion

Globally, the accumulation of fecal biomass from superabundant animals poses ecological, health, and management challenges in urban and peri-urban environments¹. Untreated droppings can contribute to nutrient loading in waterways, support pathogen transmission, and generate considerable sanitation costs ^14–18^. Bioconversion of animal feces using insect larvae offers a promising nature-based strategy to sustainably mitigate these impacts, especially because they have been widely used in bioconverting food and agricultural waste ^23,24^. Our study expands the scope beyond these applications to assess a circular system that involves two valorization pathways. First, we assess the ability of black soldier fly larvae to degrade and develop on wild bird feces (conversion of waste to insect protein), a waste stream largely overlooked in insect bioconversion research. Second, we evaluate the potential of frass derived from goose feces to fertilize plants (conversion of waste to organic fertilizer), thereby providing a complementary pathway for nutrient recycling and circular resource use. The larvae successfully completed their development across all treatments, including mixtures (50% goose feces and 50% control) as well as 100% goose feces diet. Sterilization of the media negatively affected larval performance, suggesting that fresh media with its native microbiota provides greater nutritional benefits for the larvae. Notably, frass derived from goose feces supported the growth of duckweed (a high-protein plant) at rates that exceeded those recorded for fresh feces as well as nutrient-rich Hoagland’s solution, revealing that insect digestion improves the quality of the bird feces as fertilizers. We discuss the potential of BSF larvae for sustainable waste management of animal feces.

Our records of large flocks in various urban parks of Montreal during the pre-migration period of the fall are concordant with the widespread nature of the species in Canada ^42^. Canada goose is one of the most abundant waterbird species in Canada, and their large numbers cause a nuisance to green spaces ^20^. The average number of fresh fecal deposition (deposited the same day) was 0.54 (0.15-1.18) feces m^-1^, which is within the range of estimates of fecal deposition rate recorded in western Alaska (∼0.5-2.5 feces m^-2^ day^-1^) ^36^. As expected, we documented a positive correlation between the size of the flock and the number of droppings recorded in the park. This suggests that the species spends a substantial amount of time foraging in urban parks ^43^, which is consistent with its biological needs to build fat reserves for autumn migration ^42^. Such accumulation of droppings can influence soil nutrients and plant productivity ^36^. It is unclear whether the natural degradation of this fecal material is slower in urban compared to rural environments where coprophagous communities are typically more diverse ^44^. The accumulation of a large number of droppings of the Canada goose highlights that effective artificial management involving in-situ collection and subsequent processing is necessary to remediate this environmental issue.

The growth rate of BSF in our control treatment was similar to previous reports ^45–47^. Reduced performance of BSF reared on animal excrements compared to high-quality diets such as Gainesville or chicken feed has been widely documented ^48^. Gainesville-fed larvae were 31–70% heavier than those fed poultry, swine, or dairy manure, with poultry manure yielding better growth than mammalian manure ^49^; a pattern confirmed with different types of BSF strains ^50^. Given that Canada geese primarily consume grasses, shoots, and aquatic plants, their droppings likely contain low protein and limited nutrients essential for larval optimal development. This interpretation is consistent with our lower consumption and waste reduction efficiency estimates for larvae reared exclusively on goose feces. Nonetheless, BSF larvae successfully reached adulthood, albeit more slowly, and their growth rate on goose feces was similar to that reported for chicken manure ^51^. Interestingly, a 50% mixture of Gainesville and goose feces produced growth comparable to the control, suggesting that goose manure can be effectively combined with nutrient-rich substrates to improve larval performance and yield.

The impact of autoclaving on BSF growth depended on the substrate: autoclaving the Gainesville diet enhanced growth, likely by breaking down complex molecules into simpler, bioavailable nutrients, whereas autoclaving 100% goose manure reduced growth, probably due to the loss of beneficial microorganisms and vitamins supporting larval metabolism ^52^. Similarly, a reduction in nutritional value and delayed BSF development have been reported following the drying and rehydration of manures ^53^, while autoclaving was shown to decrease survival and adult body size in sepsid dung flies, highlighting the role of microbial communities in supporting insect growth ^54^. Dry-matter consumption ranged from 54.4% in the F100 sterile treatment to 64.0% in the control, consistent with other studies reporting 35% for self-fermented coconut endosperm ^55^, 43.27% for growing-pig manure (Shao), 46.4% and 3.2% for 20% and 100% spent mushroom substrate (SMS) ^56^, and 48.41% for fermented maize straw ^57^. Waste reduction index values were lower than those reported for fast-food waste (7.86 g DM d□^1^,) and chicken feed (6.11 g), and pig manure slurry mixed with silage grass or solids (1.08–1.71 g) ^58^, as well as for 20% SMS (7 g) and 100% SMS (0.5 g) ^56^). Collectively, these findings confirm that WRI declines sharply as substrate quality decreases, driven by low nutrient content and high fibre levels.

While BSF was able to complete its life cycle in both mixed and pure goose feces, development and adult performance declined with increasing goose feces content. Concordant with other studies ^47,59^, development time was shorter in the higher quality diet than in animal manure. Body mass of BSF adults was considerably smaller in the 100% Canada geese feces than in the control and mixture, consistent with larval growth results, likely due to lower mass accumulation during larval development in nutrient-poor goose feces. The optimal C/N ratio for BSF growth ranges between 18:1 and 16:1 ^60^, whereas goose manure typically shows much higher ratios (47:1–49:1) ^61^, compared to dairy manure (21.8:1) and other herbivore waste (13.9:1–59.2:1) ^62^. Although we did not measure the nutrient composition of goose feces, its herbivorous origin suggests suboptimal protein:carbohydrate and C/N ratios for BSF larvae, a hypothesis that warrants further analysis. The 50% feces treatment, however, did not reduce adult body mass substantially, suggesting that goose feces can be combined with more nutritious substrates to sustain high performance. Adult lifespan (9 [3–12] d) was slightly shorter than those reported by other studies (e.g., 12.3-14.1 d ^63^ and 11-15 d ^64^) and was reduced under the 100% but not the 50% feces treatment. This aligns with prior evidence of a positive correlation between adult body mass and lifespan ^35^, indicating that larval nutrition carries over to adulthood. Since BSF adults rely on larval reserves to meet metabolic demands, the nutrient-poor 100% goose manure diet likely limited protein accumulation, reducing longevity. Similar effects were observed in *Telostylinus angusticollis*, where larvae on protein-restricted diets produced shorter-lived adults ^65^. Flies reared on mixed diets showed lifespans comparable to the control, and no significant effects of sex or autoclaving were detected.

In addition to producing insect protein, we showed that Canada goose feces can be used either fresh or after BSF digestion as a fertilizer to grow duckweed. Notably, we found that the growth performance of duckweed was faster in the frass than in the fresh feces, indicating that the BSF larvae improved the nutritional value of the goose feces for the duckweed. It is likely that the digestion of the goose feces by BSF larvae made key nutrients more accessible to the plant, thus enhancing nutritional quality and supporting duckweed growth ^33^. This hypothesis is strengthened by our findings on root length, exhibiting shorter roots in the frass treatment, which is a typical trait reported in laboratory and field studies in high-nutrient environments ^66,67^. Further research is needed to analyze the nutrient composition before and after BSF digestion to determine which elements are most influenced. We found a peak duckweed growth at 10 g·L^-1^, which is relatively higher than the optimal concentration reported in ^35^ showing a peak duckweed growth between 4 and 6 g·L^-1^ of BSF frass based on Gainesville diet. Our results on the benefits of Canada goose feces add more evidence to the existing literature on the positive effects of composting animal manure on soil chemical properties, growth, and yield of various crops including maize (poultry manure) ^68^, lettuce (cow manure) ^69^, and tomato (pig manure) ^70^. Even if manure involves some sanitary issues, our results on sterilization via autoclaving, which should have eliminated bacterial activity before the larval processing, showed small effects on the benefits of frass on duckweed growth rates, suggesting that nutritional value of goose feces and frass for duckweed was not largely altered after autoclaving. This suggests that the physico-chemical changes induced by BSF larvae resulted in similar nutrient availability regardless of sterilization by autoclaving. Interestingly, duckweed in both fresh feces and frass grew faster than in the high-nutrient Hoagland’s E-media, indicating that these media can serve as a great source of essential nutrients, and thereby represent a cost-effective fertilizer alternative for a wide range of crops, especially those that showed promising positive response to BSF frass such as kale ^71^, tomatoes ^72^, maize ^31^, and French beans ^73^. Future studies should explore the nutritional composition and microbial communities of Canada goose feces before and after BSF decomposition to understand the abiotic and biotic changes caused by BSF activity and assess its potential as a safe fertilizer for different edible plants.

In conclusion, our research underscores the potential of black soldier fly (BSF) larvae as effective biodegradation agents for managing Canada goose feces, an animal waste that has been historically ignored. These findings highlight the need to explore other coprophagous insects with similar or complementary bioconversion abilities, which could be integrated into fecal waste management systems to enhance efficiency and resource recovery ^74^. Large-scale implementation of such systems should involve careful planning of all components—including collection, transportation, breeding, rearing, processing, and product utilization—while ensuring safety, biosecurity, and sustainable energy use ^75^. These systems could be implemented at a small scale and tailored to urban parks, farms, or remote areas where conventional waste treatment is logistically difficult or costly. The modular nature of insect-based composting allows for flexible implementation, particularly in regions facing waste management bottlenecks or fertilizer shortages. Such integrations not only align with circular economy principles by transforming waste into valuable products but also address key sustainable development goals, such as responsible consumption and production (SDG 12) and climate action (SDG 13). Future studies should explore the safety of using BSF larvae reared on Canada goose feces as a supplement in animal feed, as well as the effectiveness of BSF frass as a plant fertilizer. Additionally, applying this bioconversion process to other waste systems, such as livestock manure or other wildlife feces, could further expand its impact, offering a scalable and eco-friendly solution to diverse organic waste challenges.

## Methods

### Black Soldier Fly Breeding

We established our black soldier fly (BSF) colony in July 2023 using 500 larvae (6–12 mm in length) obtained from a regional supplier (©The Dragon Lair, Ottawa, Canada). The larvae were reared using Gainesville (GV) diet ^76^ in 1 L rectangular plastic containers maintained at 30°C, 12:12 light-dark photoperiod, and 80% humidity. The GV diet is made of 50% wheat bran, 30% alfalfa, and 20% corn, with water added in an amount equal to the mass of the dry ingredients. Once the larvae pupated, they were collected and placed in a container within a 30 × 30 × 30 cm mesh cage (©BugDorm, Taichung, Taiwan). After emerging, adult flies were provided with water, sugar, and corrugated cardboard strips for oviposition. The colony was maintained with a population of approximately 300–500 adults, housed across four cages. To facilitate mating, adults were exposed to LED lights with a specific spectral composition designed to enhance BSF mating (Adsol^TM^ Advanced Light Solutions, Lasalle, Canada). Egg clutches were collected daily and transferred to new containers, where they were reared on fresh GV diet. Rearing was performed at these conditions for one year until the beginning of the experiment.

### Canada goose field survey

To determine whether the number of Canada geese is correlated with the number of droppings (feces), we sampled 12 urban green spaces in Southern Quebec (N=8) and Ontario (N=4), Canada (Fig. 1b). Each site was visited once during autumn (September and October) 2024 at between 11:00-15:00 (Table S1). In each site, we first counted the number of the Canada geese. Then, we counted the number of fresh feces pellets along a 100-m transect across the foraging area of the Canada goose flock. Fresh feces were readily identifiable using the dark green color and wetness of the pellet. To determine the percentage of water content in the droppings, 12 fresh droppings were collected. The fresh weight was measured using a calibrated electronic balance. Subsequently, the droppings were dried in an oven at 45°C for a duration of 96 hours.

### Bioconversion of geese feces using BSF

To test the performance of BSF larvae as bioconverters of Canada goose feces (Experiment 1), we first manually collected about 5 kg of fresh feces using plastic gloves and placed them in 19 L-plastic buckets from Angrignon Park (Montreal) on 18 September 2024. Within two hours, we transported the collected samples to the Concordia University (Loyola campus) and refrigerated them at 5 (°C). Our experimental design included three diet treatments and two sterilisation treatments (autoclaved and non-autoclaved). The three diets had varying levels of fecal content: (1) F0: 0% goose poop and 100% Gainesville (GV) diet (control), (2) F50: 50% wet GV diet + 50% wet goose feces, and (3) F100: 100% wet goose feces. The feces sterilization involved placing both GV diet and geese feces in the autoclave (fast exhaust cycle with 25 min at the sterilization phase; Fig. 1c). Each of the six treatments was replicated three times. In each replicate, we placed 100 six-day-old BSF larvae in 200 g of diet. Then, we repeated the three previously described treatments using the autoclaved substrates. We assessed larval growth by measuring fresh body weight every 2 days. To evaluate larval development, we recorded the dates and number of individuals that progressed to the pupal and adult stages daily. Additionally, we estimated adult lifespan by placing 3 males and 3 females from each replicate in individual crystal vials and recording the number of days until death. To compare adult body size across treatments, we dried the adults used for the lifespan measurements at 45°C for 48 hours and measured their dry body mass. To calculate the dry-matter reduction (hereafter referred to as the consumption rate), the following equation was applied:

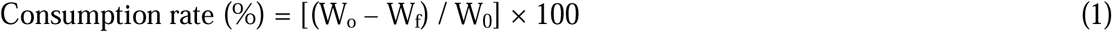

where W_o_ denotes the initial dry weight (g) of the substrate and W_f_ the remaining (frass) dry weight (g) after larval bioconversion. The waste-reduction index (WRI) was then derived by normalising the absolute reduction to the duration of exposure:

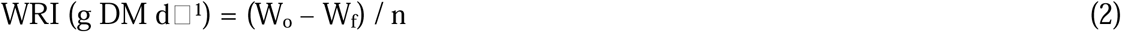

with n representing the number of rearing days, counted until at least ten larvae per replicate had reached the sixth instar or younger.

### Potential of BSF frass based on geese feces as fertilizer

*Lemna minor* was sourced from the University of Waterloo (CPCC 490) and shipped in four 25 mL tubes. Upon arrival at Concordia University’s Loyola campus, the duckweed was cultivated in a single 50 L aquarium containing sterile Hoagland’s E-media (See detailed recipe at Table S2, ^77^. The aquarium was positioned under five LED light strips, providing an intensity of 30 μmol/m²s with a 16:8 light-dark photoperiod at a constant temperature of 20 °C. The duckweed was acclimated under these conditions for six months before the experiment, with growth rates monitored and Hoagland’s E-media replenished as needed.

To evaluate the potential of Canada goose feces (both fresh and processed by BSF [frass]) in supporting duckweed growth, we conducted an experiment assessing the growth performance of common duckweed (*Lemna minor*) across 14 fertilization treatments (Experiment 2, Fig. 1d). These included two controls—distilled water [N=1] and high-nutrient Hoagland’s E-media (Table S12) [N=1] (Jewell et al., 2023)—and a full-factorial combination of feces processing by BSF (feces or frass [N=2]), autoclaving (non-autoclaved or autoclaved [N=2]), and three concentrations (2.5, 5, and 10 g·L□^1^ [N=3]). These concentrations were based on a previous study that assessed the benefit of BSF frass on duckweed growth ^35^. Duckweed was cultivated in 25 mL transparent cups in a growth chamber at 24°C and 16:8 light-dark photoperiod cycle with a light intensity of 70 μmol/m²s. For the BSF-processed goose feces treatments, we autoclaved the frass used from 100% goose feces treatment (autoclaved and non-autoclaved) of Experiment 1 (N=2). For the fresh goose feces treatments, we used both the refrigerated fresh feces and an identical media that was placed in the autoclave (N=2). Each of the 14 treatments was replicated five times. For each replicate, three randomly selected duckweed colonies (groups of 1–3 attached fronds) were placed in 25 mL cups (Diameter: 4.5 cm; Height: 3 cm). Water levels were checked daily and readjusted to maintain the initial volume (25 mL). To assess duckweed growth, photographs of each cup were taken every four days. Sampling continued until fronds in at least one treatment completely covered the water surface in all replicates. The total number of live green fronds per cup per day was quantified using ImageJ ^78^.

To assess the effect of different media on duckweed root development, we conducted a controlled experiment in which individual duckweed plants were placed in separate tubes containing one of five treatments: Hoagland’s solution (control), feces at 5 and 10 g□L□^1^, and frass at 5 and 10 g□L□^1^. Each treatment was replicated three times. The initial root length in each replicate was 1 cm. After two weeks of growth under standardized conditions (growth chamber at 24°C and 16:8 light-dark photoperiod), root length was measured for 15 randomly selected individuals per treatment using an electronic caliper with a precision of 0.01 mm.

### Statistical analyses

Statistical analyses for each variable were performed using R 4.4.3 ^79^. Assessed variables for each analysis are presented in Appendix 1. Details of the data analysis are presented in Supplementary material.

### Field survey

We tested for the correlation between the number of Canada goose (log-transformed) and the number of fresh droppings (log-transformed) using a simple linear model. Our data on the number of droppings did not exhibit spatial autocorrelation, as determined by the Moran test (Moran’s I: standard deviate = 0.392, P = 0.347) performed with the moran.test function from the spdep package ^80^.

### BSF life history

ANOVAs and generalized linear models (GLMs) were applied to assess differences in life history traits (survival rate, consumption rate, waste reduction index, adult body mass, and lifespan) across diet and autoclaving treatments. Response variables expressed as percentages were transformed using the arc-sine transformation (*asin* function) prior to analysis. The appropriate probability distribution for each response variable was determined using the simulateResiduals() function from the DHARMa package ^81^ to ensure model adequacy and goodness-of-fit (Appendix 2). The effect of two-way (Diet and autoclaving) and three-way (Diet, autoclaving, and sex) interactions were checked. The most suitable model for each experiment was selected based on the lowest AICc score. Tukey’s post hoc pairwise comparisons between levels of the categorical explanatory variables were conducted by estimating marginal means and computing contrasts using the emmeans() and contrast() functions from the emmeans 1.10.4 R package ^82^.

### Duckweed growth

To determine whether growth rate of duckweed differed significantly across our treatments, we performed stepwise model selection using likelihood ratio tests on non-linear mixed-effects models fitted with the nlme function from the nlme package and a logistic function (SSlogis). These models included log-transformed counts of duckweed fronds as a response variable, replicate as a random effect, as well as media type, concentration, and autoclave as predictor variables. The individual, additive, and interactive effects of these predictors were tested on the asymptote (Asym), infliction point of the logistic curve (xmid), and maximum growth rate (scal) (Table S11).

### Duckweed root growth

To assess differences in root length among treatments, we performed a one-way ANOVA (diet as a factor) followed by a Tukey’s HSD post hoc test to identify which treatments differed significantly.

## Data availability

All data used in this study were obtained from open-access sources. Data analyses were conducted using R version 4.4.3. The raw data supporting the findings of this study are publicly available on FigShare at https://doi.org/10.6084/m9.figshare.30279640.v1.

## Code availability

The R scripts used to perform the statistical analyses and generate the figures in this study are not publicly available but may be provided to qualified researchers upon reasonable request to the corresponding author.

## Acknowledgements

We are grateful to Yasmine Hoballah, Jordi Vilanova i Broto, and all students who helped in the different phases of the study.

## Author contributions

C.L.-M., L.R.A.-V., and R.K. conceived the study and designed the experimental framework. C.L.-M. performed the field surveys, laboratory experiments, data analyses, and prepared the figures. H.M. contributed to data processing, statistical validation, and manuscript editing. L.R.A.-V. and R.K. provided supervision, conceptual guidance, and critical review of the manuscript. C.L.-M. drafted the original manuscript, and all authors contributed to revisions and approved the final version.

