## Supplementary for "Using insects for sustainable waste management of superabundant animals"

### Supplementary Figures

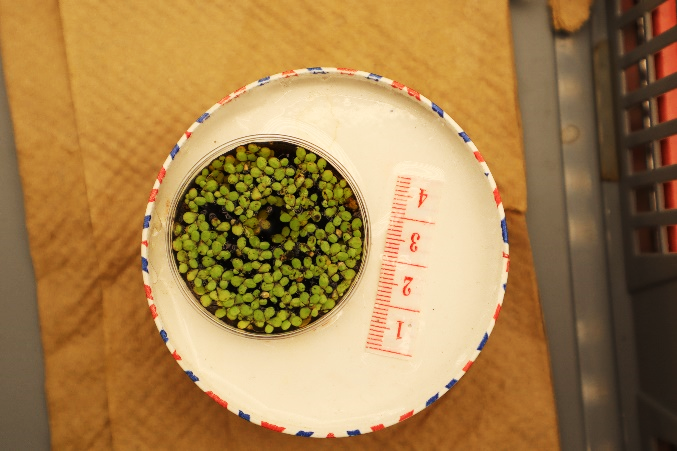

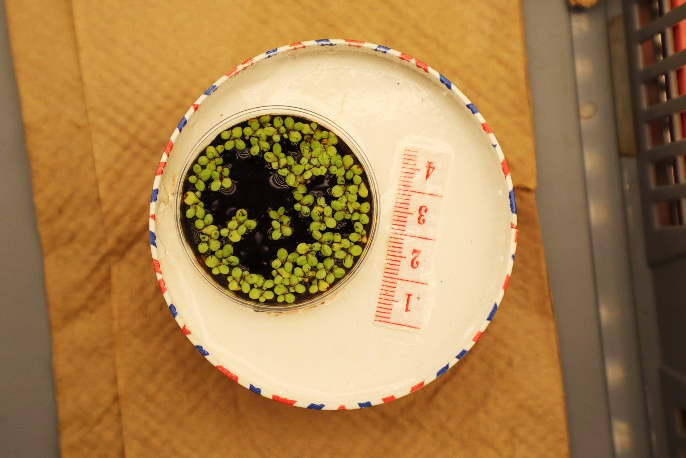

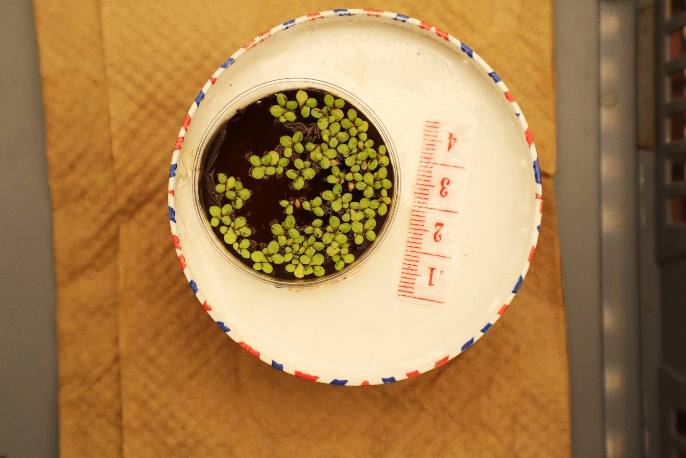

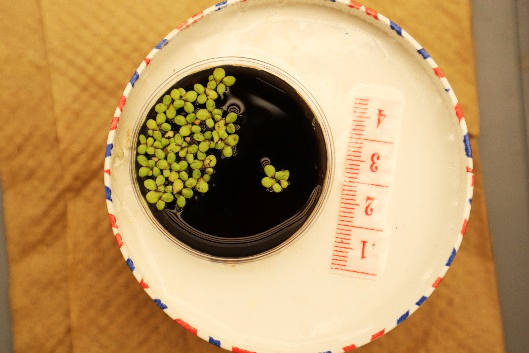

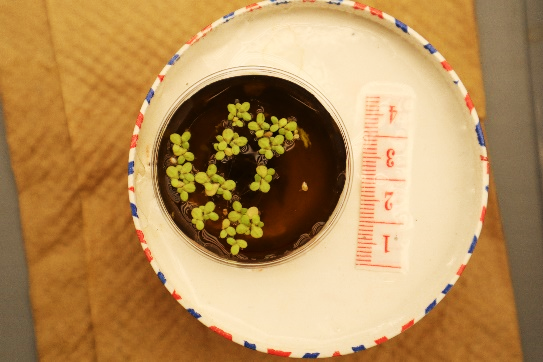

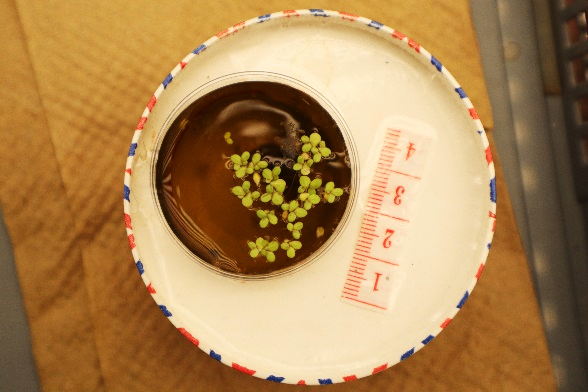

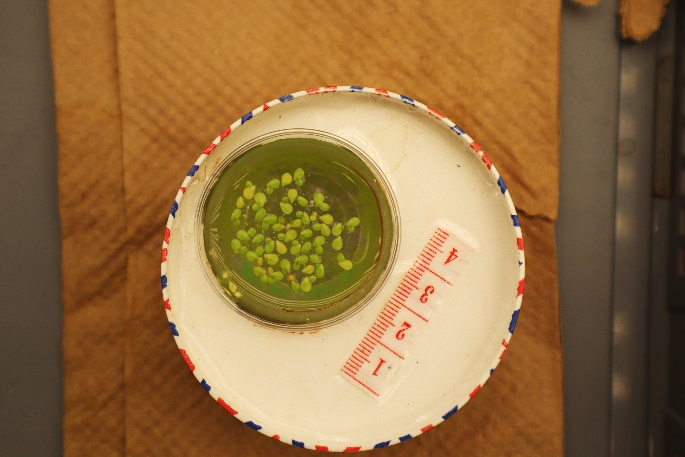

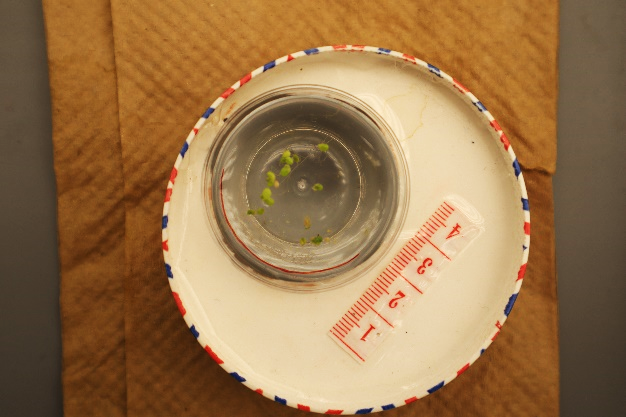

**A**

**B**

**C**

**D**

**E**

**F**

**G**

**H**

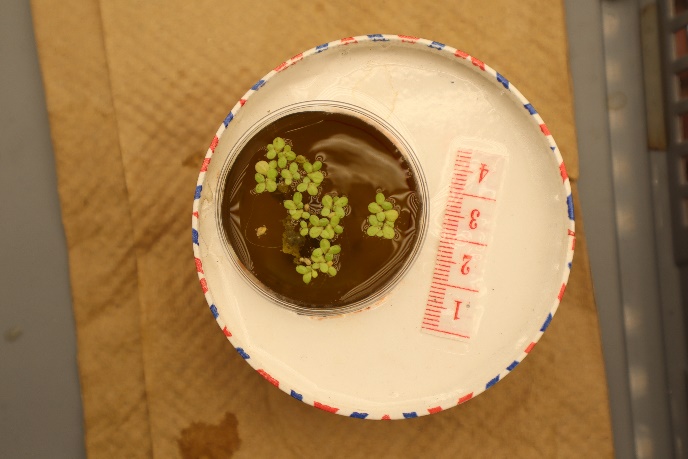

**I**

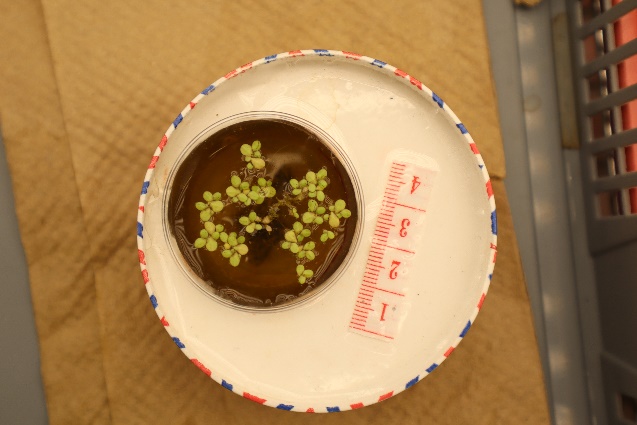

**J**

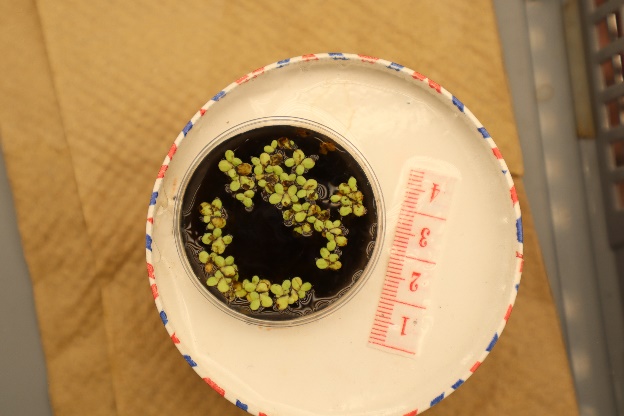

**K**

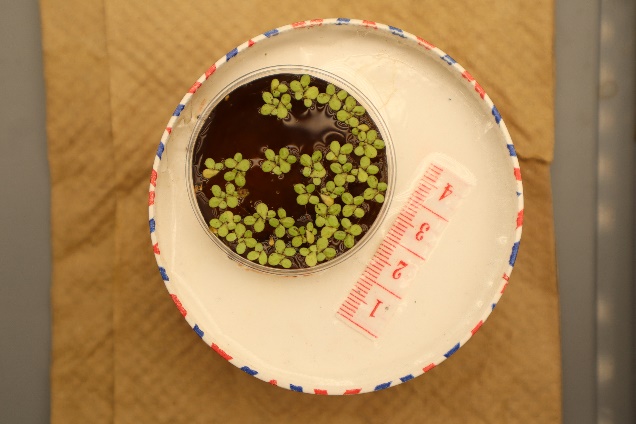

**L**

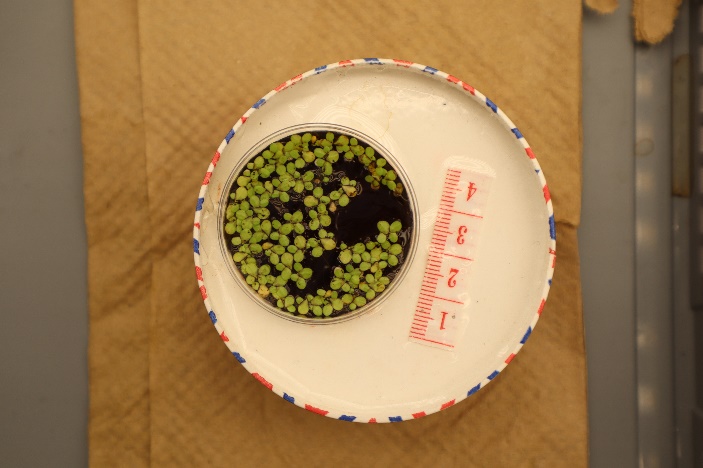

**M**

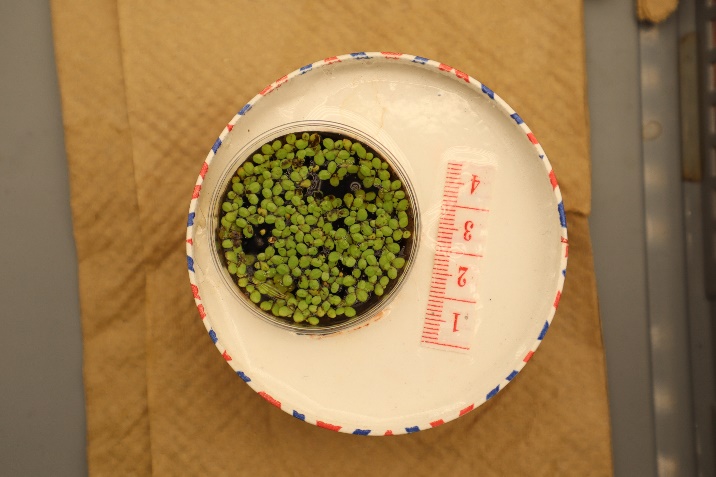

**N**

Figure S1. Duckweed response to frass and feces treatments | Growth of duckweed at 28 days in different media: (A) distilled water (control), (B) Hoagland’s solution (control), sterilised feces at (C) 2.5 g·L⁻¹, (D) 5 g·L⁻¹, (E) 10 g·L⁻¹; sterilised BSF frass of Canada goose at (F) 2.5 g·L⁻¹, (G) 5 g·L⁻¹, (H) 10 g·L⁻¹; raw feces at (I) 2.5 g·L⁻¹, (J) 5 g·L⁻¹, (K) 10 g·L⁻¹; and raw BSF frass of Canada goose at (L) 2.5 g·L⁻¹, (M) 5 g·L⁻¹, (N) 10 g·L⁻¹.

### Supplementary Tables

Table S1. Survey data: goose and dropping counts | Canada goose and fresh feces counts at 12 urban green spaces in Southern Quebec and Ontario, Canada, during September–October 2024.

| **Date** | **Site** | **Latitude** | **Longitude** | **Number birds** | **Number droppings** |
| --- | --- | --- | --- | --- | --- |
| Sep. 14 | Parc La Chine Canal East (QC) | 45.463403 | -73.599334 | 50 | 49 |
| Sep. 15 | Parc Angrignon (QC) | 45.445521 | -73.604787 | 73 | 82 |
| Sep. 18 | Parc Angrignon (QC) | 45.445522 | -73.604788 | 138 | 98 |
| Sep. 20 | Parc Jardin Botanique (QC) | 45.559052 | -73.558791 | 40 | 44 |
| Sep. 24 | Parc La Chine Canal West (QC) | 45.443266 | -73.641135 | 18 | 23 |
| Sep. 24 | Parc La Chine Canal Mid (QC) | 45.453879 | -73.622745 | 22 | 18 |
| Sep. 27 | Parc René Levesque (QC) | 45.429077 | -73.684288 | 121 | 83 |
| Oct. 05 | Parc Pierre Elliott Trudeau (QC) | 45.474619 | -73.671296 | 251 | 118 |
| Oct. 17 | Parlement Ottawa (ON) | 45.424005 | -75.699032 | 41 | 31 |
| Oct. 17 | Park des Commissaires (ON) | 45.397513 | -75.706322 | 18 | 15 |
| Oct. 19 | Burlington Waterfront Park (ON) | 43.319499 | -79.800767 | 89 | 73 |
| Oct. 20 | Kingston Confederation Park (ON) | 44.229671 | -76.478858 | 15 | 25 |

*Coordinates system: WGS 84 (EPSG:4326).*

Table S2. BSF larval growth across feces diets | Marginal growth rates (mg d⁻¹ ± 95%CI) during the first week and mean maximum body mass (mg ± 95%CI) recorded of black soldier fly larvae (*Hermetia illucens*) reared in different diets with increasing proportions of Canada goose feces.

| **Diet** | **Autoclave** | **Growth rate ± 95%CI** | **Maximum mass ± 95%CI** |
| --- | --- | --- | --- |
| F0 | No | 33.5 [31.1-35.8] | 225 [213-237] |
| F0 | Yes | 39.1 [36.8-41.4] | 259 [242-275] |
| F50 | No | 39.0 [36.7-41.4] | 240 [227-252] |
| F50 | Yes | 41.0 [38.7-43.4] | 253 [238-268] |
| F100 | No | 22.4 [20.1-24.7] | 215 [205-224] |
| F100 | Yes | 20.1 [17.8-22.5] | 191 [181-201] |

F0: 0% goose poop and 100% Gainesville (GV) diet (control), F50: 50% wet GV diet + 50% wet goose feces, and F100: 100% wet goose feces.

Table S3. Two-way ANOVA of consumption rates | Two-way ANOVA (Type II) for consumption rate, testing the effects of diet, sterilisation (Autoclave), and their interaction.

| **Term** | **Df** | **F-value** | **p-value** |
| --- | --- | --- | --- |
| Diet | 2 | 42.91 | < 0.001 |
| Autoclave | 1 | 1060 | 0.0069 |
| Diet:Autoclave | 2 | 0.89 | 0.4349 |
| Residuals | 12 |  |  |

Table S4. Post-hoc Tukey comparisons of diet consumption rates | Mean differences in consumption rate (%) between diet treatments, estimated from Tukey-adjusted pairwise comparisons (n = 18).

| **Contrast** | **Mean difference (mg) ± 95%CI** | **p-value** |
| --- | --- | --- |
| F50 – F0 | -1.67 [-4.03; 0.70] | 0.1860 |
| F100 – F0 | -7.79 [-10.15; -5.43] | < 0.001 |
| F100 – F50 | -6.12 [-8.48; -3.76] | < 0.001 |

F0: 0% goose poop and 100% Gainesville (GV) diet (control), F50: 50% wet GV diet + 50% wet goose feces, and F100: 100% wet goose feces.

Table S5. Two-way ANOVA of Waste Reduction Index (WRI) | Two-way ANOVA (Type II) for WRI, testing the effects of diet, sterilisation (Autoclave), and their interaction.

| **Term** | **Df** | **F-value** | **p-value** |
| --- | --- | --- | --- |
| Diet | 2 | 125.87 | < 0.001 |
| Autoclave | 1 | 0.066 | 0.802 |
| Diet:Autoclave | 2 | 0.514 | 0.611 |
| Residuals | 12 |  |  |

Table S6. Tukey comparisons of Waste Reduction Index (WRI) by diet | Mean differences in WRI (g DM d⁻¹) between diet treatments, estimated from Tukey-adjusted pairwise comparisons (n = 18).

| **Contrast** | **Mean difference (mg) ± 95%CI** | **p-value** |
| --- | --- | --- |
| F50 – F0 | 0.307 [-0.18; 0.79] | 0.252 |
| F100 – F0 | -2.343 [-2.83; -1.86] | < 0.001 |
| F100 - F50 | -2.65[-3.14; -2.16] | < 0.001 |

Table S7. Summary statistics of the generalized linear model (GLM) for BSF development time across different diets | Final quasi-Poisson GLM coefficients (two-way interaction model) of black soldier fly (*Hermetia illucens*) reared in different diets with increasing proportions of Canada goose feces.

| **Term** | **Estimate ± SE** | **t-value** | **p-value** |
| --- | --- | --- | --- |
| Intercept | 3.6549 ± 0.0247 | 147.8 | < 0.001 |
| Diet_F50 | –0.1157 ± 0.0366 | –3.16 | 0.0016 |
| Diet_F100 | 0.2897 ± 0.0354 | 8.19 | < 0.001 |
| Autoclave[Yes] | 0.0872 ± 0.0353 | 2.47 | 0.0136 |
| SexMale | 0.1979 ± 0.0334 | 5.93 | < 0.001 |
| Diet_F50:Autoclave [Yes] | –0.1305 ± 0.0519 | -2.51 | 0.0121 |
| Diet_F100:Autoclave [Yes] | –0.1284 ± 0.0498 | -2.58 | 0.0100 |
| Diet_F50:Sex[Male] | –0.1295 ± 0.0503 | -2.57 | 0.0102 |
| Diet_F100:Sex[Male] | –0.3731 ± 0.0499 | -7.47 | < 0.001 |

F0 [Intercept]: 0% goose poop and 100% Gainesville (GV) diet (control), F50: 50% wet GV diet + 50% wet goose feces, and F100: 100% wet goose feces.

Table S8. Post-hoc Tukey comparisons of BSF adult body mass by diet. | Tukey HSD pairwise differences among diet levels (pooled across sterilisation and sex) of black soldier fly (*Hermetia illucens*) reared in different diets with increasing proportions of Canada goose feces.

| **Contrast** | **Mean difference (mg) ± 95%CI** | **p-value** |
| --- | --- | --- |
| F50 – F0 | -3.98 [-5.7; -2.27] | < 0.001 |
| F100 – F0 | -16.23 [-17.94; -14.51] | < 0.001 |
| F100 – F50 | -12.24 [-13.96; -10.53] | < 0.001 |

F0: 0% goose poop and 100% Gainesville (GV) diet (control), F50: 50% wet GV diet + 50% wet goose feces, and F100: 100% wet goose feces.

Table S9. Logistic regression of the growth rate of duckweed in different media types | The model tests the additive effect of media type, concentration, and sterilisation (autoclave [Yes or No]). Medium is shown as type_concentration_autoclave. The baseline contrast in the intercept is distilled water. Concentrations are 2.5, 5, and 10 g/L.

|  | Value | Std.Error | DF | t-value | p-value |
| --- | --- | --- | --- | --- | --- |
| Asym.(Intercept) | 2.973 | 0.061 | 416 | 48.535 | <0.0001 |
| Asym.Medium_Frass_10_No | 3.035 | 0.130 | 416 | 23.363 | <0.0001 |
| Asym.Medium_Frass_10_Yes | 3.137 | 0.146 | 416 | 21.457 | <0.0001 |
| Asym.Medium_Frass_2.5_No | 2.312 | 0.114 | 416 | 20.316 | <0.0001 |
| Asym.Medium_Frass_2.5_Yes | 2.476 | 0.119 | 416 | 20.785 | <0.0001 |
| Asym.Medium_Frass_5_No | 2.499 | 0.117 | 416 | 21.427 | <0.0001 |
| Asym.Medium_Frass_5_Yes | 2.410 | 0.120 | 416 | 20.079 | <0.0001 |
| Asym.Medium_Hoagland_5_No | 1.602 | 0.101 | 416 | 15.861 | <0.0001 |
| Asym.Medium_Feces_10_No | 1.796 | 0.101 | 416 | 17.813 | <0.0001 |
| Asym.Medium_Feces_10_Yes | 1.774 | 0.113 | 416 | 15.660 | <0.0001 |
| Asym.Medium_Feces_2.5_No | 1.758 | 0.111 | 416 | 15.807 | <0.0001 |
| Asym.Medium_Feces_2.5_Yes | 1.677 | 0.111 | 416 | 15.161 | <0.0001 |
| Asym.Medium_Feces_5_No | 1.825 | 0.105 | 416 | 17.464 | <0.0001 |
| Asym.Medium_Feces_5_Yes | 2.239 | 0.120 | 416 | 18.676 | <0.0001 |
| xmid.(Intercept) | -18.449 | 3.696 | 416 | -4.992 | <0.0001 |
| xmid.Medium_Frass_10_No | 23.049 | 3.781 | 416 | 6.096 | <0.0001 |
| xmid.Medium_Frass_10_Yes | 25.504 | 3.802 | 416 | 6.709 | <0.0001 |
| xmid.Medium_Frass_2.5_No | 20.266 | 3.775 | 416 | 5.368 | <0.0001 |
| xmid.Medium_Frass_2.5_Yes | 21.435 | 3.779 | 416 | 5.672 | <0.0001 |
| xmid.Medium_Frass_5_No | 20.812 | 3.774 | 416 | 5.514 | <0.0001 |
| xmid.Medium_Frass_5_Yes | 21.719 | 3.782 | 416 | 5.743 | <0.0001 |
| xmid.Medium_Hoagland_5_No | 16.368 | 3.788 | 416 | 4.321 | <0.0001 |
| xmid.Medium_Feces_10_No | 16.119 | 3.780 | 416 | 4.264 | <0.0001 |
| xmid.Medium_Feces_10_Yes | 20.669 | 3.793 | 416 | 5.450 | <0.0001 |
| xmid.Medium_Feces_2.5_No | 20.073 | 3.790 | 416 | 5.296 | <0.0001 |
| xmid.Medium_Feces_2.5_Yes | 19.969 | 3.792 | 416 | 5.266 | <0.0001 |
| xmid.Medium_Feces_5_No | 17.707 | 3.780 | 416 | 4.684 | <0.0001 |
| xmid.Medium_Feces_5_Yes | 21.866 | 3.787 | 416 | 5.774 | <0.0001 |
| scal | 9.763 | 0.373 | 416 | 26.179 | <0.0001 |

Asym: Asymptote, xmid: time at the inflection point, and scal: scale parameter inversely related to growth rate

Table S10. Logistic regression (including interactions) of the growth rate of duckweed in different media types | The model tests the additive and significant interactive effects of media type, concentration, and sterilisation (autoclave [Yes or No]). The baseline contrast in the intercept is frass treatment at a concentration of 2.5 g/L.

|  | Value | Std.Error | DF | t-value | p-value |
| --- | --- | --- | --- | --- | --- |
| Asym.(Intercept) | 5.770 | 0.186 | 357 | 30.939 | <0.0001 |
| Asym.Medium_Feces | -1.282 | 0.195 | 357 | -6.563 | <0.0001 |
| Asym.Conc5 | 0.088 | 0.205 | 357 | 0.428 | 0.6691 |
| Asym.Conc10 | 0.058 | 0.192 | 357 | 0.304 | 0.7611 |
| Asym.Autoclave_Yes | 0.401 | 0.142 | 357 | 2.832 | 0.0049 |
| Asym.Medium_Feces:Conc5 | 0.179 | 0.225 | 357 | 0.794 | 0.4276 |
| Asym.Medium_Feces:Conc10 | 0.209 | 0.217 | 357 | 0.964 | 0.3358 |
| Asym.Medium_Feces:Autoclave_Yes | -0.220 | 0.110 | 357 | -1.991 | 0.0473 |
| Asym.Conc5:Autoclave_Yes | -0.247 | 0.116 | 357 | -2.134 | 0.0335 |
| Asym.Conc10:Autoclave_Yes | -0.253 | 0.115 | 357 | -2.207 | 0.0279 |
| Asym.Medium_Feces:Conc5:Autoclave_Yes | 0.361 | 0.153 | 357 | 2.364 | 0.0186 |
| Asym.Medium_Feces:Conc10:Autoclave_Yes | -0.053 | 0.151 | 357 | -0.349 | 0.7273 |
| xmid.(Intercept) | 3.478 | 0.734 | 357 | 4.739 | <0.0001 |
| xmid.MediumFeces | -3.910 | 0.585 | 357 | -6.682 | <0.0001 |
| xmid.Conc5 | 0.015 | 0.712 | 357 | 0.021 | 0.983 |
| xmid.Conc10 | 0.214 | 0.654 | 357 | 0.327 | 0.7435 |
| xmid.Autoclave_Yes | 2.500 | 0.506 | 357 | 4.944 | <0.0001 |
| scal.(Intercept) | 12.643 | 1.140 | 357 | 11.087 | <0.0001 |
| scal.MediumFeces | -3.624 | 1.452 | 357 | -2.496 | 0.013 |
| scal.Conc5 | -0.545 | 1.295 | 357 | -0.421 | 0.6741 |
| scal.Conc10 | -3.385 | 1.149 | 357 | -2.947 | 0.0034 |
| scal.Autoclave_Yes | 0.449 | 0.734 | 357 | 0.612 | 0.5409 |
| scal.Medium_Feces:Conc5 | 0.607 | 1.877 | 357 | 0.323 | 0.7466 |
| scal.Medium_Feces:Conc10 | 2.478 | 1.740 | 357 | 1.424 | 0.1552 |

Asym: Asymptote, xmid: time at the inflection point, and scal: scale parameter inversely related to growth rate

Table S11. Duckweed growth curve parameters | Parameters of the growth curves for duckweed (*Lemna minor*) in different media.

| **Parameter** | **Description** |
| --- | --- |
| Asym | a numeric parameter representing the asymptote. |
| xmid | a numeric parameter representing the x value at the inflection point of the curve. The value of SSlogis will be Asym/2 at xmid. |
| scal | a numeric scale parameter on the input axis. |

Table S12. Hoagland’s E-media composition | Recipe for Hoagland’s E Medium used in high nutrient concentrations. The pH was set to 5.8 before autoclaving the media.

| **Chemical compound** | **High concentration (mg**·**L^-1^)** |
| --- | --- |
| MgSO_4_ | 12.300 |
| Ca(NO₃)₂·4H₂O | 27.140 |
| KH_2_PO_4_ | 4.353 |
| KNO_3_ | 12.625 |
| H_3_BO_3_ | 71.500 |
| MnCl_2_ · 4H_2_O | 45.500 |
| ZnSO_4_ · 7 H_2_O | 5.500 |
| Na₂MoO₄·2H₂O | 2.250 |
| CuSO_4_ · 5 H_2_O | 3.500 |
| FeCl_3_ · 6 H_2_O | 0.484 |
| EDTA | 1.500 |

### Supplementary Results

Adult body mass*.* Diet sterilisation did not have a significant effect on adult body weight (F_1,103_= 0.23, P=0.633). Females were 23.2% heavier than males across treatments (F_1,103_= 19.9, P<0.0001). The significant diet-by-sex interaction (F_2,103_=3.79, P=0.02) indicates that the difference in dry weight between sexes was minimal in the 100% goose manure diet compared to the F0 and F50. The significant diet-by-autoclave interaction (F_2,103_=4.25, P=0.017) indicates that the reduction in adult dry mass was steeper in autoclaved compared to non-autoclaved diets (Fig. 4b).

Adult lifespan*.* No significant differences were found between F0 and F50 (z=1.92, P = 0.133). Males and females had similar lifespans (z=-0.178, P=0.859). Sterilising the media did not influence the lifespan of adults (z=1.14, P=0.252) (Fig. 4c).

Duckweed growth rate*.* Duckweed exhibited a higher growth potential in the frass treatments compared to both feces and control treatments (Table S9). Plants grown in distilled water showed minimal growth, as reflected in the lowest asymptote estimate (2.97 [95 % CI: 2.83–3.16] fronds), while those grown with frass at 10 g·L⁻¹ reached the highest population size (6.02 [95 % CI: 5.68–6.37] fronds). Compared to the feces treatments, increasing frass concentration led to a significantly higher asymptote and a later inflection point, indicating more sustained and productive growth over time. For example, asymptotes for frass at 2.5, 5, and 10 g·L⁻¹ were 5.32 [95 % CI: 4.99–5.49], 5.40 [95 % CI: 5.07–5.57], and 6.02 [95 % CI: 5.68–6.37] fronds, respectively. In contrast, feces treatments showed lower asymptotes: 4.65 [95 % CI: 4.50–4.80] at 2.5 g·L⁻¹ (reference level), 4.95 [95 % CI: 4.64–5.12] at 5 g·L⁻¹, and 4.74 [95 % CI: 4.59–4.91] at 10 g·L⁻¹. The inflection point (xmid) for frass at 10 g·L⁻¹ was significantly higher (4.83 days [95 % CI: 3.41–6.25]), suggesting a longer but more productive growth phase, while xmid estimates for other frass and feces treatments were not significantly different from the baseline. The Hoagland solution also supported robust growth, with an asymptote of 4.57 [95 % CI: 4.38–4.77] fronds, though still lower than that of frass at 10 g·L⁻¹. Autoclaving had no significant effect on asymptote (4.72 with autoclave vs. 4.65 without; p = 0.19), but significantly delayed the inflection point (xmid increased by 2.02 days [95 % CI: 1.23–2.81], p < 0.0001), suggesting that sterilisation slowed early growth dynamics. To test for interactions between media, concentrations, and autoclaving, we refitted the model, keeping only the frass and feces treatments. We found a significant three-way interaction between media, concentration, and autoclaving on the asymptote (Table S10), indicating that the effect of autoclaving depended on the media and concentration. For instance, autoclaving increased the asymptote in the frass treatment at 5 g·L⁻¹ and 10 g·L⁻¹, but decreased it at 2.5 g·L⁻¹; however, in the feces treatment, it decreased the asymptote at 2.5 g·L⁻¹ and 10 g·L⁻¹, but had no effect at 5 g·L⁻¹. The time at the inflection point was shorter in the absence of autoclaving (t = 4.94, P < 0.0001) regardless of the concentration.

#### Supplementary Methods

Canada goose field survey*.* For the simple linear model log-transformed droppings to log-transformed goose counts (Fig. 1), a model adequacy was evaluated with DHARMa (Appendix 2). Uniformity of scaled residuals was confirmed (exact one-sample Kolmogorov–Smirnov: D = 0.161, p = 0.865). A non-parametric dispersion test showed no over- or under-dispersion (ratio = 0.917, p = 0.984). An exact binomial outlier test identified no outliers (0 / 12 observations, p = 1). Spatial autocorrelation of droppings was assessed with Moran’s I under randomisation (k-nearest-neighbour weights, k = 3; spdep v. 1.3-1); none was detected (Moran’s I = ‑0.135, SD = ‑0.264, p = 0.604).

Larval growth rate*.* Model adequacy was evaluated with DHARMa v. 0.4.6. Residuals deviated marginally from uniformity (Kolmogorov–Smirnov D = 0.079, p = 0.003), but dispersion was appropriate (ratio = 0.98, p = 0.74) and only 10 of 540 observations were mild outliers (exact binomial p = 0.013). Effect sizes were obtained with Type-II Wald tests (car::Anova) and marginal growth rates (mg d⁻¹) and 95 % CI with emmeans v. 1.10.4.

Development time*.* An initial Poisson GLM revealed severe over-dispersion (AER::dispersiontest, α = 2.82, p < 0.001). All subsequent analyses therefore used a quasi-Poisson GLM with dispersion parameter φ = 4.63. The Sex × Diet interaction (Table S7) confirmed earlier female emergence in every diet (p < 0.05).

*Adult body mass.* A Type-II ANOVA (aov) was performed to assess differences in adult dry body mass across diet type (F0, F50, F100), sterilisation (No, Yes), and sex (Female, Male). Post-hoc pairwise comparisons were conducted using Tukey’s HSD (Table S8)*.*

Lifespan*.* We conducted a Poisson GLM since no over-dispersion was detected (dispersion parameter ≈ 0.64, DHARMa dispersion test p = 0.008 indicates slight under-dispersion). The final model retained diet as the only fixed effect (p=0.0134), sterilisation and sex were non-significant; p=0.8877 and p=0.0725, respectively. Model adequacy was confirmed with DHARMa (uniformity test: D = 0.117, p = 0.101; no outliers). All pairwise contrasts used estimated marginal means.

### Appendices

Appendix 1. Statistical models and distributions used | Statistical models and error distributions used to evaluate Canada goose survey and fitness traits of Black Soldier Flies (*Hermetia illucens*).

| **Analysis** | **Response variable** | **Statistical distribution** |
| --- | --- | --- |
| Canada goose field survey | Number of droppings | Normal for the residuals, after transforming the response variable using log transformation |
| Larval growth rate | Larval weight | Normal for the residuals |
| Development time | Development time | Quasi-Poisson (to account for over-dispersion) |
| Adult body mass | Dry weight | Normal for the residuals |
| Adult lifespan | Adult lifespan | Poisson distribution |
| Consumption rate | Dry mass reduction | Normal for the residuals |
| WRI | Waste reduction | Normal for the residuals |
| Duckweed growth rate | Frond count | Binomial |
| Root length | Root length | Normal for the residuals |

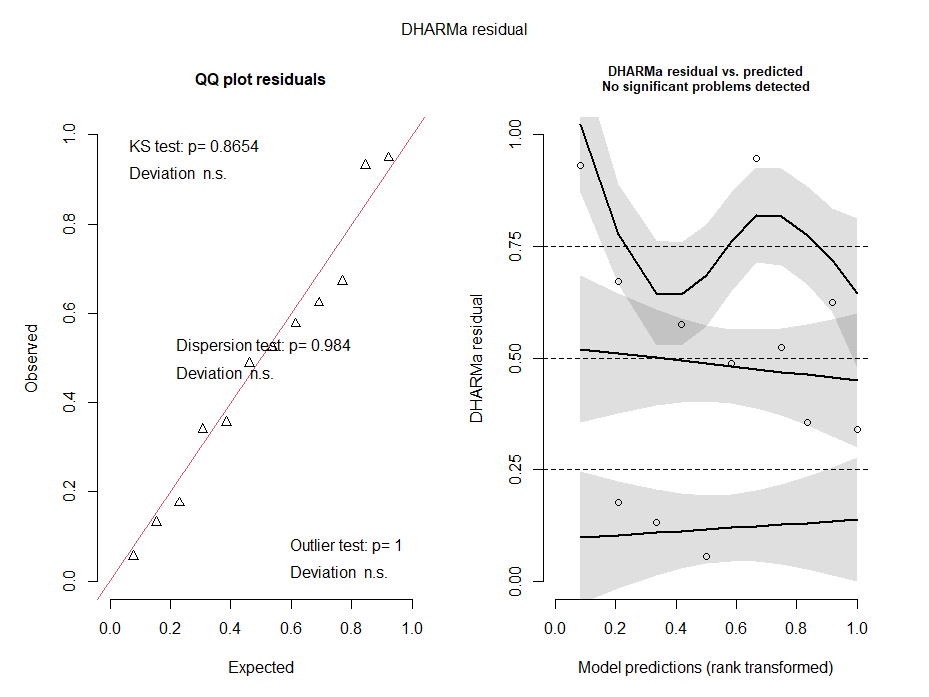

Appendix 2. Residual diagnostics of goose droppings model | DHARMa residual diagnostics for the GLM examining the relationship between the number of droppings and the abundance of Canada geese (*Branta canadensis*) in urban green spaces of southeast Canada.
